# Genetic Risk for Rheumatoid Arthritis is Associated with Increased Striatal Volume in Healthy Young Adults

**DOI:** 10.1101/519132

**Authors:** Reut Avinun, Adam Nevo, Ahmad R. Hariri

**Author notes:** Corresponding Author: Reut Avinun, Ph.D., Laboratory of NeuroGenetics, Department of Psychology & Neuroscience, Duke University, Grey Building 2020 West Main St, Ste 0030 Durham, NC 27705.

## Abstract

Rheumatoid arthritis (RA), an autoimmune disease, has recently been associated with increased striatal volume and decreased intracranial volume (ICV) in longstanding patients. As inflammation has been shown to precede the clinical diagnosis of RA and it is a known moderator of neuro- and gliogenesis, we were interested in testing whether these brain morphological changes appear before the clinical onset of disease in healthy young adult volunteers, as a function of relative genetic risk for RA. Genetic and structural MRI data were available for 516 healthy non-Hispanic Caucasian university students (275 women, mean age 19.78±1.24 years). Polygenic risk scores were computed for each individual based on a genome-wide association study of RA, so that higher scores indicated higher risk. Striatal volume (sum of caudate, putamen, and nucleus accumbens volumes) and ICV were derived for each individual from high-resolution T1-weighted images. After controlling for sex, age, genetic components of ethnicity, socioeconomic status, and depressive symptoms, we found that higher RA polygenic risk scores were associated with increased striatal volume, but not decreased ICV. Our findings suggest that increased striatal volume may be linked to processes that precede disease onset, such as inflammation, while decreased ICV may relate to disease progression.

---

Rheumatoid arthritis (RA) is an autoimmune, inflammatory disease, that is characterized by progressive articular destruction, chronic pain, and a peak age of onset in the fifth decade of life. In a recent study, Wartolowska et al.^1^ found that in comparison with healthy volunteers, patients with RA had increased striatal volume (putamen, caudate, and nucleus accumbens) and decreased intracranial volume. They further suggested that increased striatal volume may be associated specifically with the experience of chronic pain in RA patients. However, depression and inflammation are additional features of RA^2^, and these have been associated with structural alterations in various brain regions, including the striatum^3–5^. Notably, inflammation has been demonstrated to precede the onset of clinical RA^6^. Therefore, it is possible that increased striatal volume is present before the development of clinical symptoms of RA, including chronic pain.

Increasingly, genetic tools are being employed to examine disease-related processes before their clinical onset. In particular, polygenic risk scores, based on the weighted sum or average effects of individual risk alleles identified in genome-wide association studies (GWAS), have emerged as a reliable strategy to model individual differences in disease-related processes. A recent meta-analysis of GWAS^7^ encompassing 29,880 RA cases and 73,758 controls demonstrated the involvement of primary immunodeficiency genes and showed that common single nucleotide polymorphisms (SNP), outside of the major histocompatibility complex region, explain 5.5% of disease risk heritability in European Caucasians. Here, we used genetic and structural MRI data from a large sample of healthy young non-Hispanic Caucasian university students, to test if higher polygenic risk scores for RA, based on the above GWAS meta-analysis, are associated with increased striatal volume and decreased ICV, as has been reported in patients with RA, before the onset of clinical disease.

## RESULTS

Linear regression analysis indicated that RA polygenic risk scores were not significantly associated with ICV (β=-.026, SE=.033, p=.435). However, in a model examining striatal volume as the dependent variable, RA polygenic risk scores were significantly positively correlated with striatal volume (β=.089, SE=.032, p=.005; Figure 1). Of the covariates considered, sex (β=-.129, SE=.044, p=.003), age (β=-.114, SE=.030, p<.001), and ICV (β=.577, SE=.038, p<.001), were significantly correlated with striatal volume, wherein men, younger individuals, and individuals with relatively larger ICVs, were more likely to have increased striatal volume.

**Figure 1.**
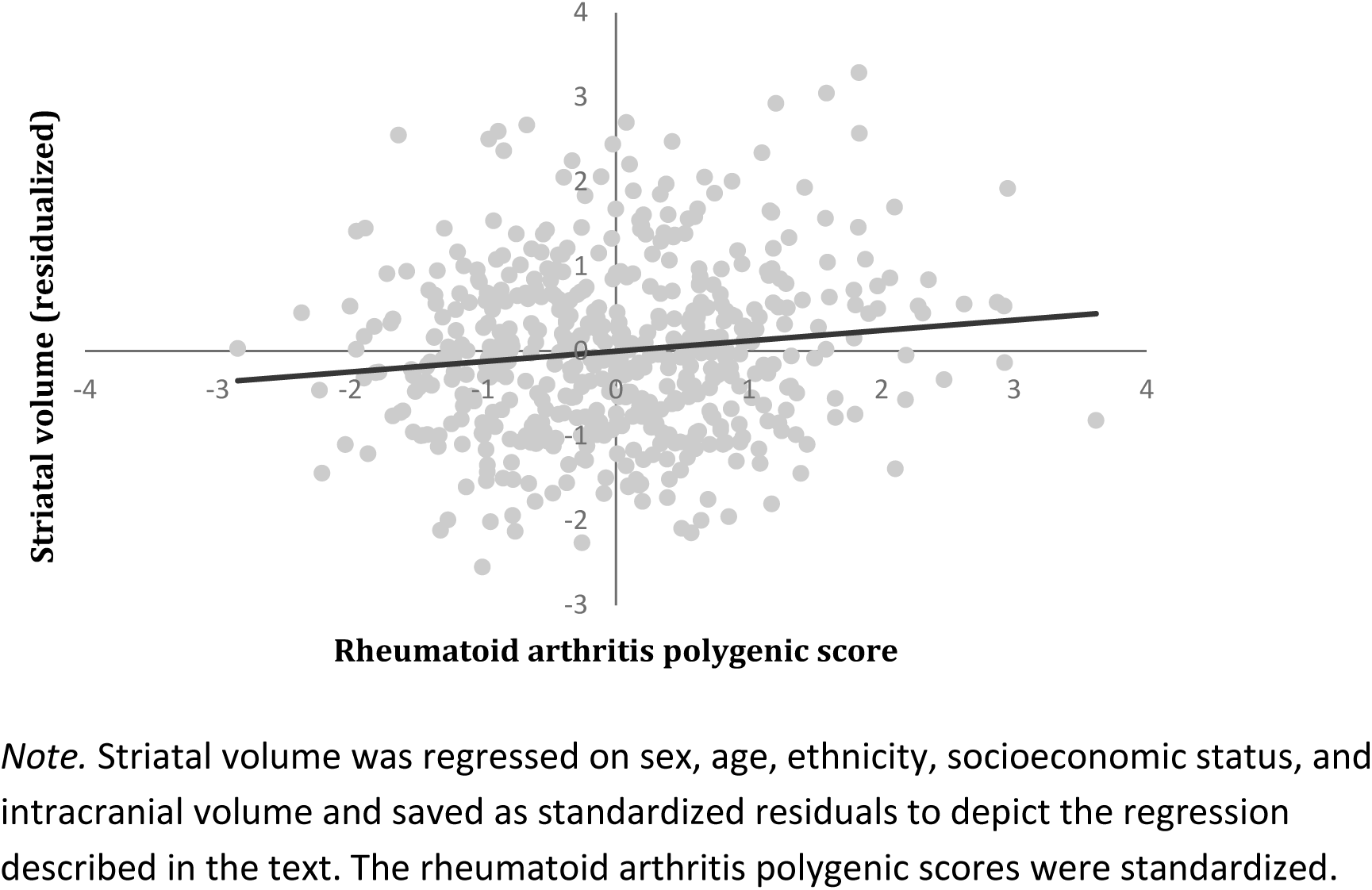
GWAS-derived rheumatoid arthritis polygenic risk scores are positively correlated with striatal volume in healthy young adults. *Note.* Striatal volume was regressed on sex, age, ethnicity, socioeconomic status, and intracranial volume and saved as standardized residuals to depict the regression described in the text. The rheumatoid arthritis polygenic scores were standardized.

As RA may be comorbid with depression^2^, and there have been links between depression and striatal volume^4,5^, we further examined the above associations with depression as a covariate. In a model controlling for depressive symptoms, which significantly and positively predicted striatal volume (N=514; β=.090, SE=.032, p=.005), RA polygenic risk scores remained significantly correlated with striatal volume (N=514; β=.091, SE=.031, p=.004).

As post-hoc analyses, we examined whether a specific striatal subregion was driving the observed effect. While the RA scores significantly and positively predicted caudate volume (β=.091, SE=.037, p=.014), and putamen volume (β=.082, SE=.037, p=.027), they were not significant predictors of nucleus accumbens volume (β=-.042, SE=.041, p=.301).

## DISCUSSION

Here, we extended the prior finding of increased striatal volume in RA^1^ by showing that higher polygenic risk scores for RA are associated with increased striatal volume in a sample of healthy, young adult volunteers. This suggests that increased striatal volume may precede the onset of clinical symptoms of RA. Post hoc analysis of striatal subregions showed that RA scores significantly and positively predicted putamen and caudate volumes, but not nucleus accumbens volume. Furthermore, these associations were independent of sex, age, and socioeconomic status as well as depressive symptoms, which have been independently associated with both striatal volume and RA^2,5^. In contrast, we did not find a significant correlation between RA polygenic risk scores and ICV unlike that observed in patients with RA ^1^. This may reflect differences between premorbid (i.e., striatal volume) and postmorbid (i.e., ICV) processes.

Wartolowska et al.^1^, hypothesized that the link between RA and increased striatal volume may be due to an effect of chronic pain^3^. The current findings in healthy young adults, suggest that alternative pathways may exist. One such pathway is inflammation. Although inflammation is typically regarded as detrimental to neurogenesis, accumulating research demonstrates that its effects are more complex, and could even enhance neurogenesis depending on various factors, including the developmental stage and the duration and location of the inflammation^8^. In addition, inflammation can promote gliogenesis^8^, which may also contribute to the observed association between higher genetic risk and larger striatal volume. Interestingly, increased striatal volume was also found in fibromyalgia patients^3^, another disease characterized by systemic inflammation^9^.

Importantly, inflammation in RA has been shown to precede clinical symptoms^6^, and genetic studies, including the GWAS meta-analysis on which we based our polygenic risk scores^7^, have demonstrated enrichment of immune related genes in risk for RA^10^. It is therefore possible that individuals with a genetic susceptibility to develop RA, have higher levels of inflammation early on, which possibly affects striatal volume before the clinical onset of RA. Further studies combining polygenic risk scores, brain imaging, and markers of inflammation are needed to evaluate these possible links.

Of course, our study is not without limitations. First, while using a polygenic risk score allowed us to examine associations with striatal volume before the onset of RA, we do not have longitudinal data into middle and late life that could enable testing associations with disease emergence. Second, our findings are limited to non-Hispanic Caucasians, and may not generalize to populations with different genetic backgrounds. Finally, while our findings are consistent with those reported by Wartolowska et al.^1^, they are novel and thus replication is necessary before any possible utility of striatal volume as a biomarker can be determined. These limitations notwithstanding, our results demonstrate that increased striatal volume, but not decreased ICV, can be found in healthy young adults at relatively higher genetic risk for RA, suggesting that increased striatal volume may be linked to processes that precede disease onset, such as inflammation, while decreased ICV may relate to disease progression.

## MATERIALS AND METHODS

### Participants

Our sample consisted of 516 non-Hispanic Caucasian participants (275 women, mean age 19.78±1.24 years) from the larger Duke neurogenetics study (DNS) for whom there was complete data on genotypes, structural MRI data, depressive symptoms, and all covariates described below. The DNS was approved by the Duke University Medical Center Institutional Review Board, and all experiments were performed in accordance with the relevant guidelines and regulations. Prior to the study, all participants provided informed consent. Notably, self-reported medication information was examined to verify that none of the participants were prescribed either disease modifying antirheumatic drugs or biological treatments. All participants were free of the following study exclusions: 1) medical diagnoses of cancer, stroke, diabetes requiring insulin treatment, chronic kidney or liver disease, or lifetime history of psychotic symptoms; 2) use of psychotropic, glucocorticoid, or hypolipidemic medication; and 3) conditions affecting cerebral blood flow and metabolism (e.g., hypertension).

Of the 516 non-Hispanic Caucasians, 114 individuals had at least one DSM-IV diagnosis as determined by structured clinical interview. Importantly, neither current nor lifetime diagnosis were an exclusion criterion, as the DNS sought to establish broad variability in multiple behavioral phenotypes related to psychopathology. However, no participants, regardless of diagnosis, were taking any psychoactive medication during or at least 14 days prior to their participation.

### Race/Ethnicity

An analysis of identity by state of whole-genome single nucleotide polymorphisms (SNPs) was performed in PLINK^11^. The first two multidimensional scaling components were used as covariates.

### Socioeconomic status (SES)

We controlled for possible SES effects using the “social ladder” instrument, which asks participants to rank themselves relative to other people in the United States (or their origin country) on a scale from 0–10, with people who are best off in terms of money, education, and respected jobs, at the top and people who are worst off at the bottom. SES ranged between 2 and 10 (M=7.35, SD=1.43 years).

### Depressive symptoms

The 20-item Center for Epidemiologic Studies Depression Scale (CES-D) was used to asses depressive symptoms in the past week^12^. All items were averaged to create a total depressive symptoms score. Depressive symptoms ranged between 0 and 43 (M=8.82, SD=7.01).

### Genotyping

DNA was isolated from saliva using Oragene DNA self-collection kits (DNA Genotek) customized for 23andMe (www.23andme.com). DNA extraction and genotyping were performed through 23andMe by the National Genetics Institute (NGI), a CLIA-certified clinical laboratory and subsidiary of Laboratory Corporation of America. One of two different Illumina arrays with custom content was used to provide genome-wide SNP data, the HumanOmniExpress (N=327) or HumanOmniExpress-24 (N=189).

### Quality control and polygenic scoring

PLINK^11^ was used to perform quality control analyses and remove SNPs or individuals based on the following criteria: missing genotype rate per individual >.10, missing rate per SNP >.10, minor allele frequency <.01, and Hardy-Weinberg equilibrium p<1e-6.

Polygenic risk scores were calculated by using PLINK^11^ and the “--score” command on the SNP-level summary statistics from a GWAS meta-analysis of RA^7^. Specifically, the summary statistics for the European Caucasian sample (14,361 RA cases and 43,923 controls) were used. Notably, as the reported effect sizes were odds ratios, they were log transformed prior to calculations. For each SNP the number of the alleles (0, 1, or 2) associated with RA was multiplied by the effect estimated in the GWAS. A polygenic risk score for each individual was an average of weighted RA-associated alleles. All matched SNPs were used regardless of effect size and significance in the original GWAS as previously recommended and shown to be effective^13,14^. Consequently, 441,939 SNPs were included in the calculation of the polygenic score. The approach described here for the calculation of the polygenic score was successfully used in previous studies e.g.,^15,16,17^.

### Structural MRI

Data were collected at the Duke-UNC Brain Imaging and Analysis Center using one of two identical research-dedicated GE MR750 3T scanners (General Electric Healthcare, Little Chalfont, United Kingdom) equipped with high-power high-duty cycle 50-mT/m gradients at 200 T/m/s slew rate, and an eight-channel head coil for parallel imaging at high bandwidth up to 1 MHz. T1-weighted images were obtained using a 3D Ax FSPGR BRAVO with the following parameters: TR = 8.148 ms; TE = 3.22 ms; 162 axial slices; flip angle, 12°; FOV, 240 mm; matrix =256×256; slice thickness = 1 mm with no gap; and total scan time = 4 min and 13 s.

To generate regional measures of brain volume, anatomical images for each subject were first skull-stripped using ANTs^18^, then submitted to Freesurfer’s (Version 5.3) recon-all with the “-noskuNstrip” option^19,20^, using an x86_64 linux cluster running Scientific Linux. The gray and white matter boundaries determined by recon-all were visually inspected using FreeSurfer QA Tools (https://surfer.nmr.mgh.harvard.edu/fswiki/QATools) and determined to be sufficiently accurate for all subjects. Volume measures for the caudate nucleus, nucleus accumbens, and putamen from each participant’s aseg.stats file were averaged across hemispheres and then summed to create a total striatal volume variable. Estimated Total Intracranial Volume (eTIV) was used to quantify intracranial volume (ICV).

### Statistical Analyses

Mplus version 7^21^ was used to conduct linear regression analyses with the following covariates: participants’ sex (coded as 0=males, 1=females), age (in the DNS 18-22 years were coded as 1-5), genetic ethnicity components, SES, and intracranial volume (when striatal volume was the dependent variable). The RA polygenic risk score, intracranial volume, and striatal volume were standardized to improve interpretability. Maximum likelihood estimation with robust standard errors, which is robust to non-normality, was used in the regression analyses. Standardized results are presented.

Author Contributions
R.A., A.N., and A.R.H. designed research; R.A. analyzed data; R.A., A.N., and A.R.H., wrote the paper.

## REFERENCES

1 Wartolowska, K. et al. Structural changes of the brain in rheumatoid arthritis. Arthritis Rheumatol 64, 371–379 (2012).

2 Kojima, M. et al. Depression, inflammation, and pain in patients with rheumatoid arthritis. Arthritis Care Res. 61, 1018–1024 (2009).

3 Schmidt-Wilcke, T. et al. Striatal grey matter increase in patients suffering from fibromyalgia–a voxel-based morphometry study. Pain 132, S109–S116 (2007).

4 Koolschijn, P. et al. Brain volume abnormalities in major depressive disorder: A meta-analysis of magnetic resonance imaging studies. Hum. Brain Mapp. 30, 3719–3735 (2009).

5 Savitz, J. et al. Activation of the kynurenine pathway is associated with striatal volume in major depressive disorder. Psychoneuroendocrinology 62, 54–58 (2015).

6 Kokkonen, H. et al. Up-regulation of cytokines and chemokines predates the onset of rheumatoid arthritis. Arthritis Rheum. 62, 383–391 (2010).

7 Okada, Y. et al. Genetics of rheumatoid arthritis contributes to biology and drug discovery. Nature 506, 376 (2014).

8 Borsini, A., Zunszain, P. A., Thuret, S. & Pariante, C. M. The role of inflammatory cytokines as key modulators of neurogenesis. Trends Neurosci. 38, 145–157 (2015).

9 Bäckryd, E., Tanum, L., Lind, A.-L., Larsson, A. & Gordh, T. Evidence of both systemic inflammation and neuroinflammation in fibromyalgia patients, as assessed by a multiplex protein panel applied to the cerebrospinal fluid and to plasma. Journal of pain research 10, 515 (2017).

10 Eyre, S. et al. High-density genetic mapping identifies new susceptibility loci for rheumatoid arthritis. Nat. Genet. 44, 1336 (2012).

11 Purcell, S. et al. PLINK: a tool set for whole-genome association and population-based linkage analyses. The American Journal of Human Genetics 81, 559–575 (2007).

12 Radloff, L. S. The CES-D scale: A self-report depression scale for research in the general population. Applied psychological measurement 1, 385–401 (1977).

13 Dudbridge, F. Power and predictive accuracy of polygenic risk scores. PLoS Genet. 9, e1003348 (2013).

14 Ware, E. B. et al. Heterogeneity in polygenic scores for common human traits. bioRxiv, 106062 (2017).

15 Domingue, B. W., Belsky, D. W., Conley, D., Harris, K. M. & Boardman, J. D. Polygenic influence on educational attainment: New evidence from the National Longitudinal Study of Adolescent to Adult Health. AERA open 1, 2332858415599972 (2015).

16 Domingue, B. W., Liu, H., Okbay, A. & Belsky, D. W. Genetic heterogeneity in depressive symptoms following the death of a spouse: Polygenic score analysis of the US Health and Retirement Study. Am. J. Psychiatry 174, 963–970 (2017).

17 Stephan, Y., Sutin, A. R., Luchetti, M., Caille, P. & Terracciano, A. Polygenic Score for Alzheimer Disease and cognition: The mediating role of personality. J. Psychiatr. Res. (2018).

18 Klein, A. et al. Evaluation of 14 nonlinear deformation algorithms applied to human brain MRI registration. Neuroimage 46, 786–802 (2009).

19 Dale, A. M., Fischl, B. & Sereno, M. I. Cortical surface-based analysis: I. Segmentation and surface reconstruction. Neuroimage 9, 179–194 (1999).

20 Fischl, B., Sereno, M. I. & Dale, A. M. Cortical surface-based analysis: II: inflation, flattening, and a surface-based coordinate system. Neuroimage 9, 195–207 (1999).

21 Muthén, L. K. & Muthén, B. O. Mplus User’s Guide. Los Angeles, CA: Muthén & Muthén (2007).

